# Spatial RNA proximities reveal a bipartite nuclear transcriptome and territories of differential density and transcription elongation rates

**DOI:** 10.1101/196147

**Authors:** Jörg Morf, Steven W. Wingett, Irene Farabella, Jonathan Cairns, Mayra Furlan-Magaril, Xin Liu, Frank F. Craig, Simon Andrews, Marc A. Marti-Renom, Peter Fraser

**Affiliations:** Laboratory of Nuclear Dynamics, The Babraham Institute, Cambridge, UK; Bioinformatics, Babraham Institute, Cambridge, UK; CNAG-CRG, Center for Genomic Regulation (CRG), Barcelona Institute of Science and Technology (BIST), Barcelona, Spain Gene Regulation, Stem Cells and Cancer Program, Center for Genomic Regulation (CRG), Barcelona Institute of Science and Technology, Barcelona, Spain; Institució Catalana de Recerca i Estudis Avançats (ICREA), Barcelona, Spain; Departamento de Genética Molecular, Instituto de Fisiología Celular, Universidad Nacional Autónoma de México, México; Sphere Fluidics Limited, Babraham Research Campus, Cambridge, UK

## Abstract

Spatial transcriptomics aims to understand how the ensemble of RNA molecules in tissues and cells is organized in 3D space. Here we introduce Proximity RNA-seq, which enriches for nascent transcripts, and identifies contact preferences for individual RNAs in cell nuclei. Proximity RNA-seq is based on massive-throughput RNA-barcoding of sub-nuclear particles in water-in-oil emulsion droplets, followed by sequencing. We show a bipartite organization of the nuclear transcriptome in which compartments of different RNA density correlate with transcript families, tissue specificity and extent of alternative splicing. Integration of proximity measurements at the DNA and RNA level identify transcriptionally active genomic regions with increased nucleic acid density and faster RNA polymerase II elongation located close to compact chromatin.

---

The spatial organization of nucleic acids in cell nuclei is critical for gene expression and ultimately cell physiology (1). However, our understanding of which transcripts are synthesized, processed and/or sequestered in the same spatial nuclear compartments lags behind progress in elucidating genome organization in space. Parts of transcriptomes have been spatially resolved in tissues (2) and single cells (3, 4). While these techniques position RNA molecules in two dimensions within the biological material they do not infer spatial associations between different RNA molecules. On the other hand pairwise probing of RNA-RNA interactions has so far been restricted to direct base-paired contacts (5) or Ångström distances between RNA ends allowing for enzymatic ligations of protein-bound molecules (6-8). We thus devised a method that determines spatial associations between pairs or groups of transcripts irrespective of the nature of interaction and not limited to direct contacts between RNA molecules (9).

Proximity RNA-seq tags RNAs contained in individual crosslinked sub-nuclear particles with unique barcodes during reverse transcription (Fig.1A, fig.S1). To tag tens of millions of particles individually and in a parallel fashion, we applied picodroplet water-in-oil emulsions. Homogenates were emulsified in oil by a simple and rapid vortexing step together with beads, each covered with uniquely barcoded random primers for reverse transcription (9). After cDNA synthesis in droplets, sequencing of all barcoded cDNA molecules enabled the reconstruction of RNA proximities in nuclei.

**Fig. 1.**
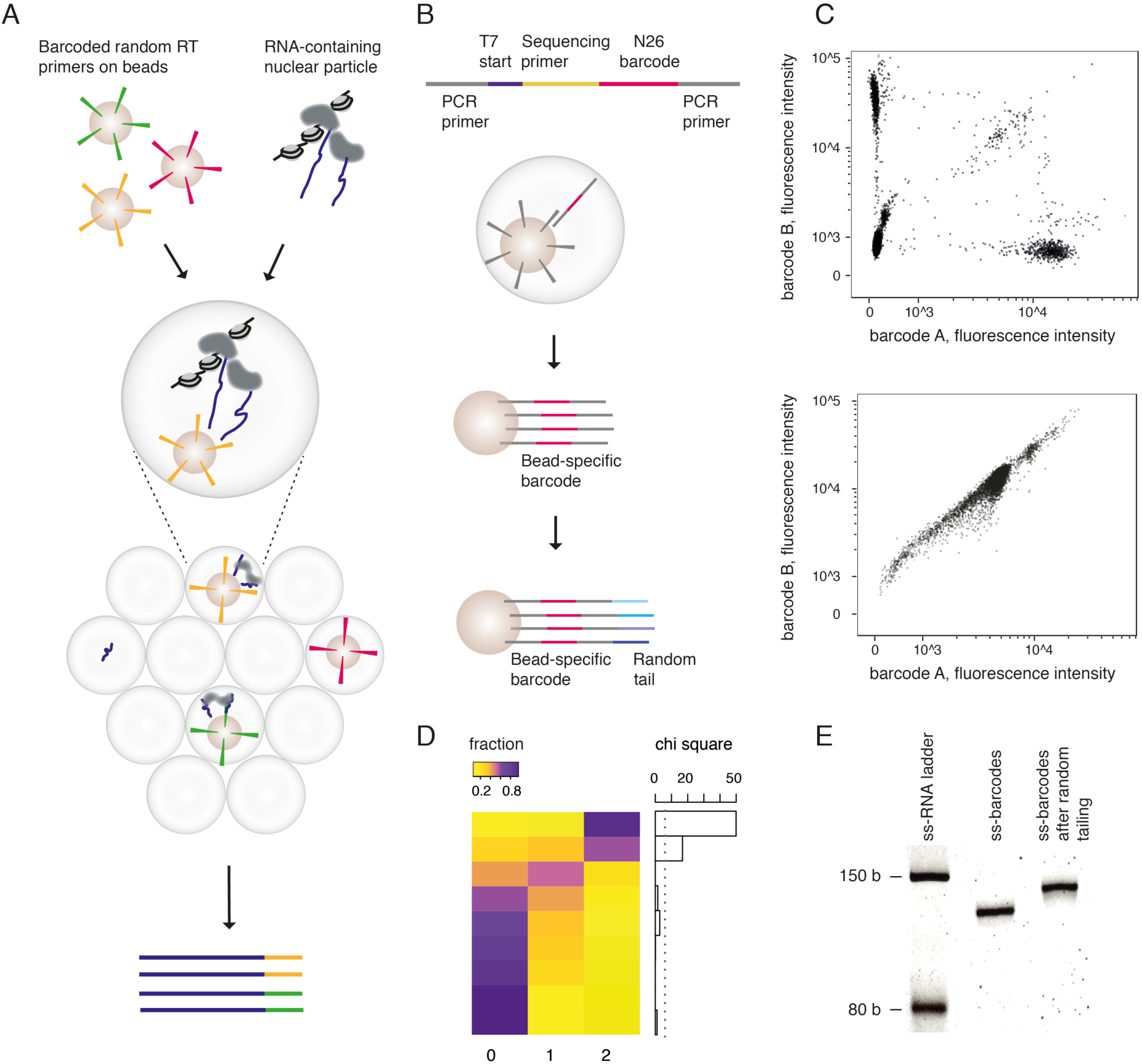
Proximity RNA-seq A) Massive-throughput barcoding by reverse transcription (RT) of RNA-containing particles in water-in-oil droplets. B) Barcoding of magnetic beads with immobilized primers and diluted random DNA templates by emulsion PCR. Barcodes on beads were end-tailed with random nucleotides to subsequently serve as RT primers. C) To control barcoding and emulsion integrity two barcodes of known sequence were amplified on beads in emulsion or solution prior to hybridization with complementary fluorescent probes and FACS analysis to count empty, single, and mixed barcoded beads. Top panel, PCR in emulsion, bottom, in solution. Axes specify fluorescence signals of hybridized probes. D) Different two-barcode experiments, rows, were ordered according to increasing fractions of non-barcoded beads (yellow: low, purple: high fraction). Fractions of beads containing no, one or both barcodes (columns) were compared to expected Poisson distributions (chi-square test, dashed line indicates p = 0.05). E) Acrylamide gel of single-stranded barcodes before (lane 1) and after (lane 2) the addition of 15 random nucleotides.

First we developed an on-bead PCR protocol in emulsion to individually barcode up to one billion beads for an experiment. Ideally, a single synthesized DNA template containing 26 random bases as barcode is amplified in a droplet containing one bead covered with immobilized primers (Fig.1B). The encapsulation of a single barcode with one bead can be approximated by diluting templates and beads sufficiently before emulsification. To optimize the yield of barcoded beads however, we chose encapsulation conditions and template amounts according to a Poisson model that aimed at 50% barcoded beads, of which around 70% were covered with copies of a single barcode (Fig.1C, D). After PCR, individual DNA barcode copies on beads were extended with random bases to generate reverse transcription primers (Fig.1B, E). Sub-nuclear particles from human neuroblastoma cells, SH-SY5Y, were obtained by mild sonication of chemically crosslinked and washed cell nuclei in a manner similar to chromosome conformation capture, Hi-C (9, 10) (fig.S2). Sequencing of such preparations resulted in RNA reads enriched for nascent transcripts as indicated by the high overall intronic read density and the characteristic overrepresentation of reads at 5’ compared to 3’ ends in introns (11) (Fig.2A, B). Reads were then grouped according to their barcodes (fig.S3, S4). Multiple reads with the same barcode and mapping to the same transcript were counted as one observation, dubbed a proxy read (9). Approximately 25% of proxy reads were co-barcoded with proxy reads mapping to other transcripts and specified spatial RNA associations (fig.S5, S6).

**Fig. 2.**
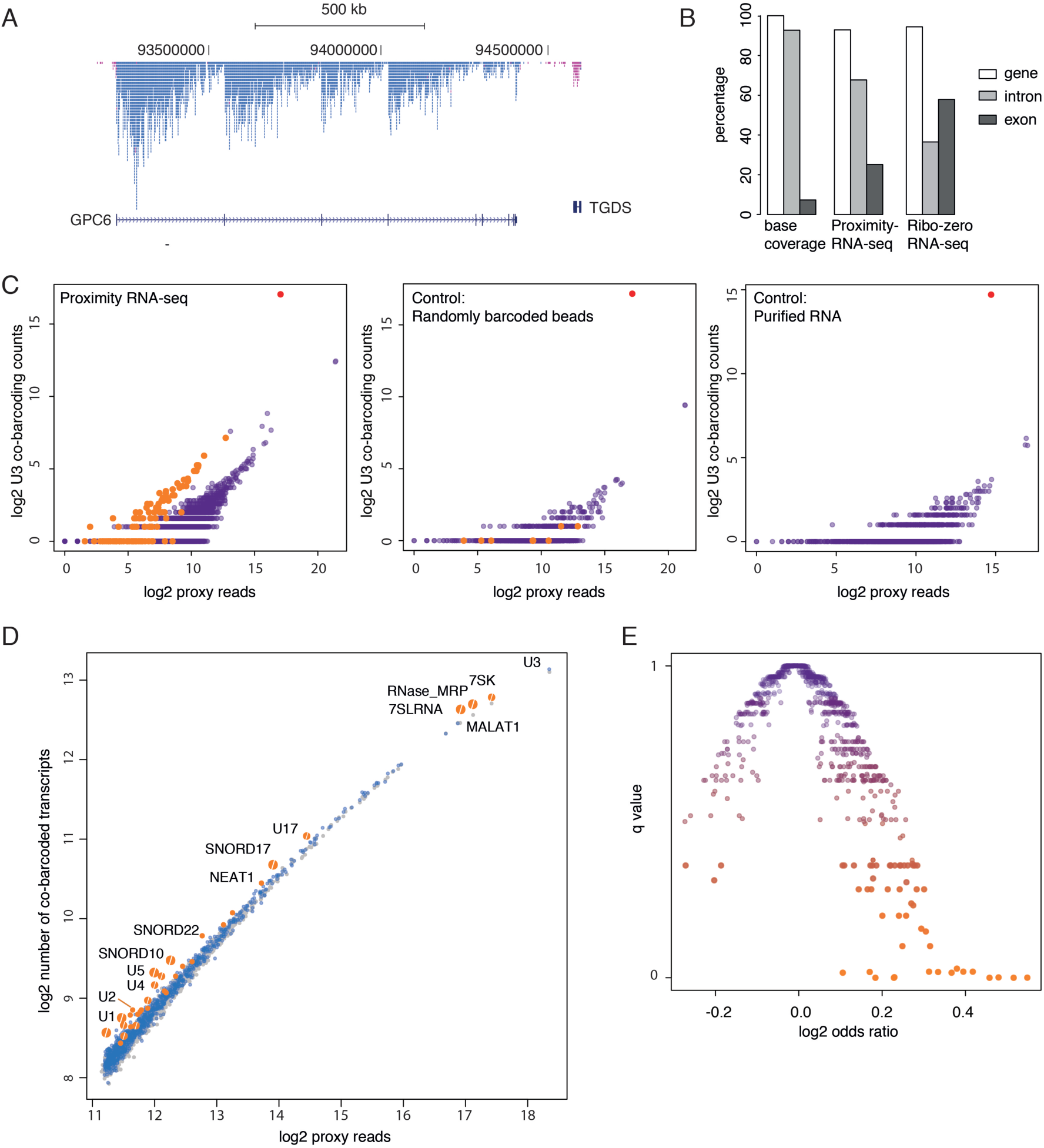
RNA co-barcoding and proximal transcriptomes. A) Proximity RNA-seq reads mapping to GPC6 (blue) and adjacent transcripts on the opposite strand (pink) illustrate both high intronic read densities and the saw-tooth read pattern along introns. B) Fractions in percentage of reads in transcript features. Proximity RNA-seq was compared to total RNA-seq after ribosomal depletion (25) and the base coverage of exons, introns and genes (set as 100), respectively. C) Counts of co-barcoding events involving U3 RNA plotted against the number of proxy reads of the other RNA. SnoRNAs are in orange, other transcripts purple, U3 red. Left panel: crosslinked sample with uniquely barcoded beads, middle: control using crosslinked sample and randomly barcoded beads, right: control with purified RNA after crosslink reversal and uniquely barcoded beads. Of note most snoRNAs are not plotted for control data, as no co-barcoding with U3 was detected. D) For the top 1000 RNAs, the number of contacted transcripts was plotted against the number of proxy reads per RNA. Grey: randomized, blue: observed, orange: RNAs with complex contactomes (p < 0.01), orange with white slash: q < 0.05 as derived in E). E) Transcripts with more complex proximal transcriptomes than expected at random were identified by Fisher’s exact test and FDR adjustment (y: q value, x: odds ratio).

To validate spatial associations we analysed co-barcoding events of the abundant small RNA U3. U3 resides in the nucleolus, a phase separation of proteins and hundreds of small non-coding RNAs (snoRNAs), which compartmentalizes ribosomal RNA synthesis, processing and ribosome subunit assembly (12). We found snoRNAs co-barcoded considerably more with U3 than expected by their abundance and compared to the non-snoRNA transcriptome (Mann-Whitney U: p 4∗10^-56^, Fig.2C). In contrast, control experiments using beads barcoded by PCR in a droplet-free solution and harbouring a high complexity of different barcodes on each bead, or crosslink-reversed RNA on uniquely barcoded beads, showed few or no co-barcoding between U3 and snoRNAs. Next, we analysed the number of unique transcripts co-barcoded, once or multiple times, with a given RNA molecule of the 1000 highest expressed transcripts. We noticed that RNAs present at higher levels, with more opportunities to encounter other transcripts, had higher co-barcoded transcript counts (Fig.2D). Notably, we identified RNAs with more complex proximal transcriptomes than expected, which included spliceosomal (U1, U2, U4 and U5), paraspeckle (NEAT1) and nucleolar (snoRNAs) transcripts (Fig.2D, E).

We then asked whether groups of transcripts preferentially associate with each other and thereby partition the nuclear transcriptome. First, to calculate the significance of spatial associations between transcripts while taking their vastly different abundances into account, we randomized pairings of proxy reads and their barcodes 100000 times to obtain simulated co-barcoding counts for pairs of transcripts. We then compared simulated against observed counts (fig.S7, S8) (9). Principal component analysis on simulation-derived p values of pairwise associations between any RNA and the top 100 connected transcripts suggested that indeed the nuclear transcriptome partitioned into two compartments (Fig.3A). Principal component 2 (PC2) separated snoRNAs as well as a set of protein-coding transcripts from the bulk of mostly protein-coding transcripts (Fig.3A-C). We dubbed the nucleolar transcript group compartment I, based on the prevalent RNA polymerase I activity for ribososomal RNA synthesis within the compartment. Accordingly, compartment II was named after RNA polymerase II, which is predominant in the nucleoplasm. We next analyzed features of RNAs grouped into 8 quantiles according to their relative nuclear position derived from PC2 values. We observed an accumulation of gene ontology terms, mostly specific to the neuronal cell type, in compartment II (quantiles 4-8) but few enriched terms in compartment I (quantiles 1-3) (fig.S9). Similarly, the number of transcripts with high tissue specificity (13, 14) increased from compartment I to II, with the exception of nucleolar quantile 1 (Fig.3D). Consistently, genes encoding compartment II transcripts were closer in the linear genome to multi-enhancer domains crucial for cell identity (super-enhancers (15, 16)) than genes whose transcripts were assigned to compartment I (fig.S9). Finally, alternative splicing of exons in protein-coding transcripts occurred less frequently in compartment I, as indicated by high exon inclusion (17). In contrast, transcripts in compartment II exhibited variable and often lower exon inclusion (Fig.3E). This suggests that broadly expressed transcripts are enriched in vicinity to nucleoli and tissue specific RNAs and transcripts that are more frequently subject to alternative splicing in compartment II, respectively.

**Fig. 3.**
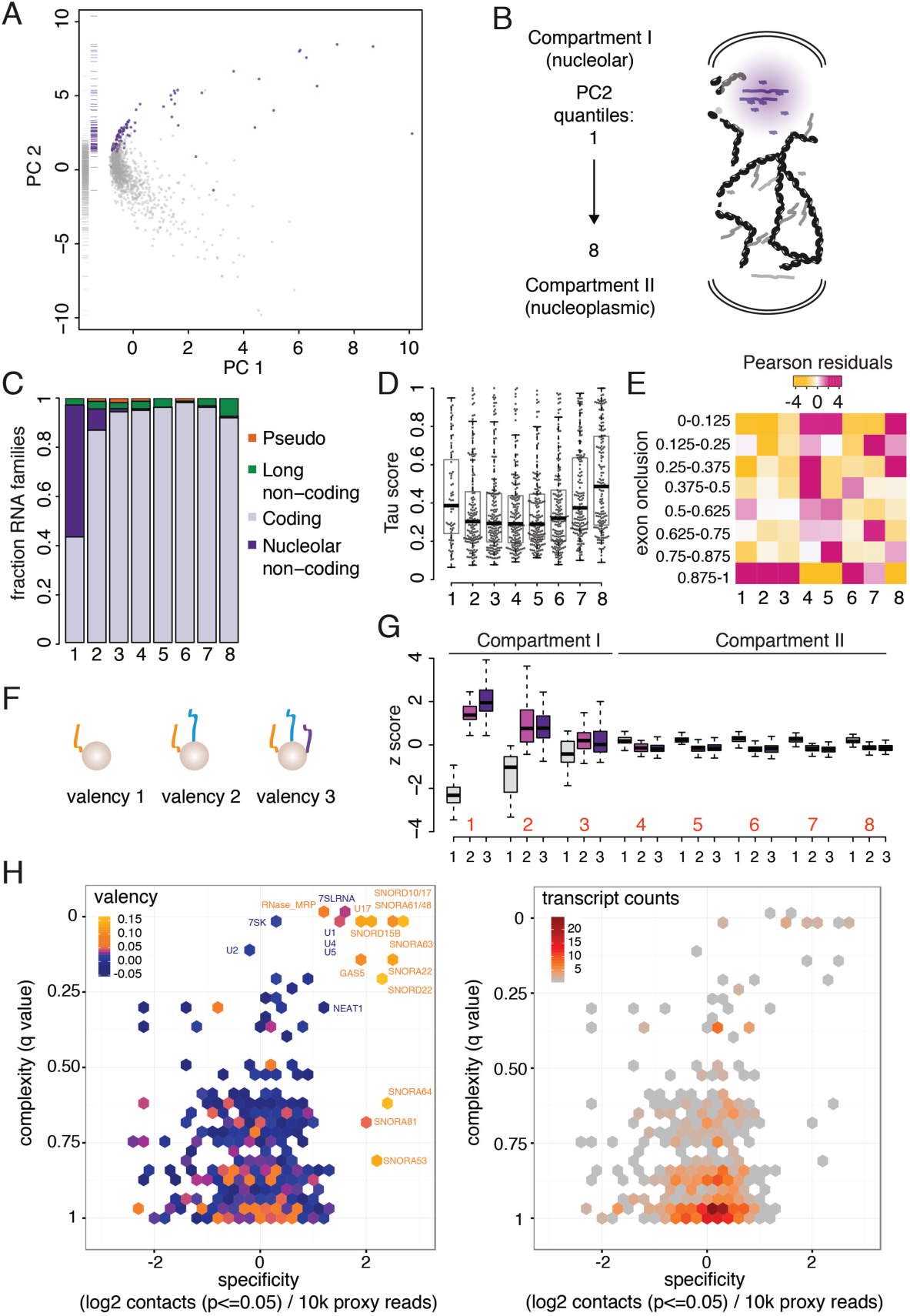
Bipartite nuclear transcriptome and RNA valency. A) PCA of transcripts based on their pairwise proximity with the top 100 connected transcripts. Purple: snoRNAs, Grey: other RNAs. B) Schematic of compartment I with transcripts in purple and compartment II with transcripts in grey. PC2 quantiles 1 - 8 are indicated. C) Fractions of RNA families along PC2 axis. D) Tissue specificity (Tau scores, 0 for broadly expressed and 1 for tissue specific (13, 14)) for each transcript in different proximity quantiles. E) Heatmap of Pearson residuals (yellow-magenta) from regression against PC2 quantiles (columns) and bins of exon inclusion scores (rows). F) RNA valency. G) Valency (x-axis: black numbers) along PC2 axis (x: red numbers). H) Transcript-specific proximal transcriptomes were described by their q values of complexity plotted against specificity, defined as the number of pairwise contacts (p <= 0.05) of a given transcript divided by its number of proxy reads. Hexagons, representing single or multiple transcripts within a given bin, were colored based on the number of transcripts per bin (right panel) or mean valency of proximal transcriptomes (left panel). For hexagon bins with multiple transcripts the highest mean valency specified the color.

Proximity RNA-seq enables the simultaneous detection of two or more proximal, co-barcoded transcripts. With the aim to derive local RNA density and/or connectivity for individual transcripts we introduced a measure describing how often a transcript was detected singly or co-barcoded with one or multiple other RNA molecules. This so-called valency of a given transcript was inferred from relative enrichments or depletions (z-score) in the number of reads mapping to the transcript in barcode groups encompassing 1, 2 and 3 transcripts, respectively, compared to the nuclear transcriptome average (Fig.3F, fig.S10, S11) (9). We found that transcripts in compartment I overall exhibited high valency, i.e. z-scores of valency “1” < 0 and valency “2” and “3” > 0, indicating the increased RNA density in nucleoli and perinucleolar regions (18). In contrast, low valency transcripts prevailed in compartment II (Fig.3G, fig.S12).

Our data shows that transcripts can be located at RNA dense or sparse spatial positions within the nucleus. Moreover, a given RNA molecule might display different complexities of proximal transcriptomes, contacting larger or smaller sets of other transcripts, and do so with variable specificity towards nearby transcripts (fig.S13). Taking into account these different specifications we combined three measures derived from Proximity RNA-seq to characterize transcripts: complexity of the proximal transcriptome (Fig.2D), specificity towards proximal transcriptomes, which we based on the number of preferential pairwise associations (p <= 0.05), and mean valency of proximal transcriptomes (pairwise contacts p <= 0.1) to estimate RNA density (Fig.3H, fig.S14). Such analysis revealed that snoRNAs had high valency and often complex proximal transcriptomes, but also maximal specificity. This further delineates compartment I as a separate structure of high RNA density consisting of a large defined set of transcripts. On the other hand, spliceosomal transcriptomes (of U1, U2, U4 and U5 RNAs), while consistently of high complexity, showed reduced specificity and valency. This is in line with the high target diversity of spliceosomes in the nucleoplasmic compartment II with lower RNA density than compartment I (Fig.3G) (19).

Two main compartments outline genome organization; open, transcriptionally active (A) and compact, lowly expressed (B) multi-megabase regions with increased contact enrichments between segments of the same compartment (10). However, it has recently been suggested that genomic segments might rather be described by a continuous than a binary compartment classification (20). To identify further sub-compartments we performed Hi-C in SH-SY5Y cells and assigned RNA valency scores to A and B segments (9) (fig.S15). We found that chromatin contacts between segments of high RNA valency exhibited stronger contact enrichments (observed/expected contacts) than pairs of low valency regions did (Fig.4A, C). We then compared contacts between segments classified either as A and B or low and high valency and found that low and high valency contact distributions resembled A to A and A to B distributions, respectively (Fig.4B). Segments without assigned valency showed strongest contact enrichments (Fig.4A), similar to contacts between B regions (Fig. 4B). Indeed, contacts between segments without valency transcripts consisted in large part of B-B pairs (Fig. 4D). Unexpectedly, low and high valency segments both corresponded to a similar high extent, 90 and 94 %, respectively, to regions assigned to the A compartment (Fig.4D). This raised the possibility that while similar in A-ness to low valency regions, high valency segments were located within or near denser chromatin to form stronger contacts (Fig.4B) (21). Accordingly, we found high valency segments of A-type more frequently to undergo inter-compartment contacts with denser B regions than low valency regions (Fig.4E). To illustrate distinct spatial distributions and local chromatin properties for different regions we three-dimensionally modelled a chromosome using Hi-C contact frequencies as spatial restraints (22). We then convoluted the ensemble of the 1000 best restraint-satisfying models in a 3D density map (9). In line with the proposed distribution of different valency segments, high valency territories in chromosome 14 were spatially closer to the B compartment (Fig.4F, fig.S16). Model-derived estimates of local accessibility for a virtual object and of chromatin contact counts per volume suggested that high valency regions were less accessible (Fig.4G, fig.S16). Finally, we asked whether nuclear regions of different valency were distinguishable by local, apparent catalytic activities. To this end, we used the decrease in read density from 5’ to 3’ of long introns measured by Proximity RNA-seq (Fig.2A) as a correlate for RNA polymerase II elongation rates (11, 23). We observed faster transcription elongation in high valency compared to low valency regions (Fig.4H, fig.S17), indicating that different nuclear territories exhibit distinct kinetic parameters of transcription. We speculate that, besides genomic sequence and chromatin determinants, sequestering or phase separation of certain protein factors might favour rapid transcription elongation. Interestingly, transcription elongation factors have been shown to associate with chromatin in an RNA-dependent manner (24), suggesting that transcripts themselves in crowded regions might accumulate elongation factors critical to speed up transcription and thereby sustain locally high RNA valency.

**Fig. 4.**
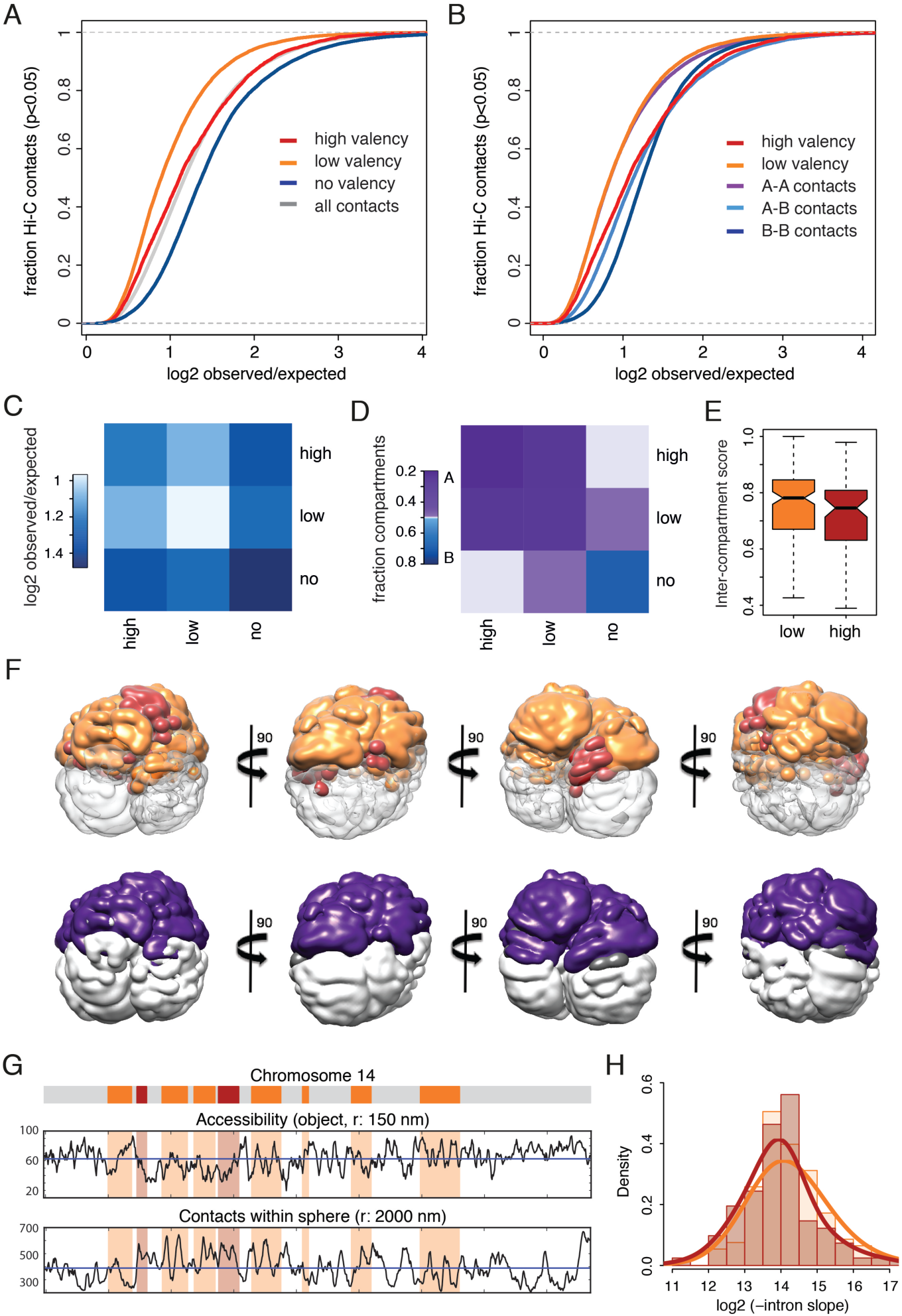
RNA valency-defined chromatin territories. A) Cumulative distributions of Hi-C contact enrichments (p <= 0.05) for high, low and no valency genomic segments (red, orange, blue). All segments are shown in grey. B) Cumulative distributions of Hi-C contact enrichments (p <= 0.05) for high and low valency segments (red, orange) in comparison to A-A, A-B and B-B contacts (purple, light blue, blue). C) Mean contact enrichments between high, low and no valency regions. D) Fractions of A (and B) regions involved in pairwise contacts between high, low and no valency regions. 0 corresponds to only A regions, 1 to exclusively B regions. E) Inter-compartment contacts of high and low valency A regions. 1 and -1 correspond to exclusively intra-(A-A) and inter-(A-B) compartment contacts, respectively (Kolmogorov-Smirnov, one-sided: p 0.02). F) Density map of chromosome 14 ensemble model. Top row: high valency regions in red, low orange, no grey. Bottom row: A, B regions, A purple, B grey, unassigned regions in dark grey. G) Local accessibility for a virtual object with radius of 150 nm and local contact density number within spherical volumes with radii 2000 nm. High valency regions in red, low orange. H) Histogram of intronic read density decays indicated faster transcription elongation in genes within high (red) than low valency (orange) regions (Kolmogorov-Smirnov: p 0.015).

In conclusion, we have established Proximity RNA-seq that portrays transcriptomes in space based on RNA proximity preferences and valency of transcripts. Such RNA measurements will enable the investigation of spatial composition of transcripts in various subcellular compartments not readily approachable with existing technologies.

## Acknowledgements

J.M. was supported by a Swiss National Science Foundation early postdoc mobility fellowship, a Human Frontier Science Program long-term fellowship and the Babraham Science Policy Committee. Work of I.F. and M.A.M-R. was partially supported by the European Research Council under the 7^th^ Framework Program FP7/2007-2013 (ERC grant agreement 609989) and the European Union’s Horizon 2020 research and innovation programme (grant agreement 676556) to M.A.M-R. M.A.M-R. was also supported by the Spanish Ministry of Economy and Competitiveness (BFU2016-75008-P), Centro de Excelencia Severo Ochoa 2013-2017 (SEV-2012-0208) and AGAUR. M.F-M. was supported by UNAM Technology Innovation and Research Support Programme (PAPIIT) num. IA201817.

We acknowledge Sphere Fluidics for their contribution of microfluidic knowhow and time and their free donation of surfactants.

We thank Babraham sequencing and FACS facilities for technical support, and Paulo Amaral and Lucas Edelman for helpful discussions.

